# Probing isn’t Causality in Molecular Models

**DOI:** 10.64898/2026.10.05.756781

**Authors:** Satya Pratik Srivastava, Sharath Krishna Chundru, Harshit Singh, Rajeev Kumar Singh

## Abstract

Linear probes establish whether a chemical variable can be recovered from a molecular representation; they do not establish how that variable contributes to subsequent predictions. We replace a concept-associated scalar component at an intermediate block and measure the change through a separately fitted down-stream outcome readout. Comparisons with random directions and architecture-matched random encoders distinguish recoverability, intervention selectivity, and a pretraining-associated selectivity advantage. In Uni-Mol, block-12 total-distortion edits predict the observed energy-change sign for 97.6% of held-out pairs, with Spearman *ρ* = 0.787. Structure-clustered intervals for the primary direction and encoder contrasts exclude zero. A stricter 20-direction sensitivity analysis does not survive correction across six cells. A post-review analysis finds substantial concept– outcome alignment; intervention effects persist after removal of the same-block outcome direction, but this does not establish that subsequent encoder computation is necessary. In MolCLR, pretraining improves dipole recovery by 0.080 *R*^2^, whereas concept-minus-random sign-accuracy contrasts are 0.029 and −0.015 at the two tested blocks. Their paired uncertainty is unresolved, so they do not establish an absence of selective effects. We propose an operational evidence ladder whose conclusions are conditional on the concept, intervention, control family, and fitted readout.

## 1 Introduction

Molecular representation learning provides features for property prediction, but predictive performance alone does not explain how internal states support a prediction. Linear probes fit a predictor of a chemical descriptor to a frozen representation. Strong held-out performance establishes recoverability, rather than necessarily establishing use of that descriptor by the encoder (Belinkov, 2022; Ravichander et al., 2021). High-dimensional features can expose variables that are incidental to the computation being studied.

We ask whether a recovered direction supports a selective intervention effect in a specified encoder– readout system. We replace a concept-associated component at an intermediate block and measure the resulting change after the remaining computation with a separately fitted outcome readout. This interchange intervention relates to causal abstraction, mediation, and activation patching (Geiger et al., 2021; Vig et al., 2020; Meng et al., 2022; Conmy et al., 2023). The intervention changes an internal state; it does not establish that the direction is unique, that it completely represents the chemical concept, or that a native property head uses it.

Our results make three contributions. First, Uni-Mol distortion edits show positive direction-specific and pretraining-associated contrasts in the primary analysis, with their scope qualified by stricter sensitivity controls. Second, total-distortion probe recovery decreases with depth while the tested intervention selectivity increases; this is a descriptive dissociation, not evidence that later blocks perform more concept-specific computation. Third, MolCLR illustrates the need to report the uncertainty of the concept-minus-random contrast rather than use a power calculation to declare an intervention null. The two systems differ in architecture, modality, training, concept, outcome, and sample size; they are distinct case studies rather than controlled experimental arms.

## 2 Related work

Probe research distinguishes information recoverable by a fitted predictor from information used by the analyzed model (Belinkov, 2022). Control tasks assess what a probe can learn from incidental features (Hewitt and Liang, 2019), and task-irrelevant properties can remain decodable (Ravichander et al., 2021). Amnesic probing tests behavioral changes after removing information (Elazar et al., 2021). Causal abstraction evaluates correspondences through interchange interventions (Geiger et al., 2021); mediation and circuit-discovery methods localize influential states or components (Vig et al., 2020; Meng et al., 2022; Conmy et al., 2023). Activation-patching conclusions also depend on intervention construction and evaluation metrics (Zhang and Nanda, 2024). Our contribution transfers these distinctions to molecular encoders with chemically matched pairs and explicit direction and encoder controls.

## 3 Method

### 3.1 Targeted interchange intervention

At encoder block *ℓ*, a ridge probe fitted to concept labels defines the unit direction in raw activation coordinates,

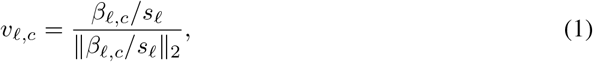

where *β*_*ℓ,c*_ is the coefficient in standardized coordinates and *s*_*ℓ*_ is the training-set feature scale. For recipient *r* and donor *d*, the edit is

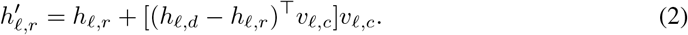

The unedited and edited states pass through the same remaining computation and outcome head. Concept and outcome heads are fitted separately, with validation used to select ridge regularization and training data used for the final fit. Separate fitting does not imply statistically independent or orthogonal directions.

### 3.2 Outcome orientation, signs, and contrasts

Let *ŷ*_*r*_ and 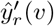 denote unedited and edited predictions for the recipient. The predicted effect and observed outcome difference are

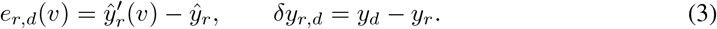

The frozen primary sign-accuracy implementation is

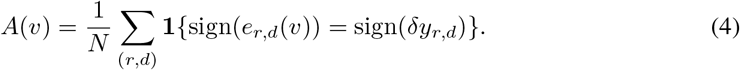

It uses exact floating-point signs, with sign(0) = 0, and no tie exclusions: a zero predicted effect is incorrect for a nonzero outcome difference, and two exact zeros agree. The post-review outcome-projection analysis instead excludes |*δy*| ≤ 10^−12^; its saved pair-level data contain no such exclusions in any of the six cells. No threshold is imposed on predicted effects. Sensitivity to near-zero predicted effects remains a numerical limitation.

For QM7-X, the optimized geometry is the recipient and a displaced conformation is the donor; the outcome is displaced-minus-optimized energy. All accepted outcome changes have the expected positive sign. Consequently, high sign accuracy there measures the frequency of an upward predicted change and should be read alongside effect–outcome Spearman correlation, effect magnitude, and direction controls. For QM9, the orientation is the saved donor-minus-recipient dipole difference. Random-direction accuracy need not equal 0.5.

For the primary random-direction set *R* and random-encoder set *Q*, define

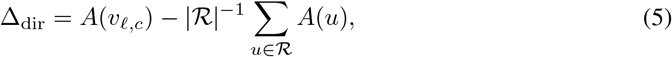

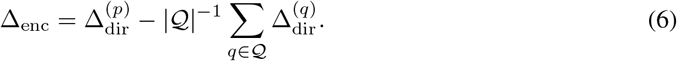

The first measures selectivity within an encoder; the second compares that selectivity between pretrained and random encoders. Five primary random directions are orthogonalized to the concept direction and to previously drawn directions, and use their own donor–recipient projections. A separate sensitivity family uses 20 exactly magnitude-matched directions. These control families are not interchangeable.

### 3.3 Systems and statistical scope

Uni-Mol uses the official 47.3M-parameter checkpoint (Zhou et al., 2023) on QM7-X (Hoja et al., 2021). Optimized structures are split into 2,708 training, 349 validation, and 311 test structures; displaced conformations stay with their optimized structure. Total and bond distortion are the concepts, and relative PBE0+MBD energy is the outcome. There are 31,100 held-out interventions. Edits occur after blocks 4, 8, and 12, and the readout is evaluated after block 15. Thus edits at block 12 traverse three further blocks before the measured readout, irrespective of any later blocks in the encoder.

MolCLR uses the official 2.21M-parameter five-block GIN (Wang et al., 2022), with 1,072 held-out same-formula, fingerprint-matched QM9 pairs (Ramakrishnan et al., 2014). A Mulliken-derived classical dipole is the concept and the DFT dipole is the outcome. The uniform edit shifts every node by the same vector so the mean-pooled state receives the donor projection. Edits follow blocks 2 and 4, and the readout follows block 5.

The frozen Uni-Mol analysis uses 2,000 bootstrap draws of optimized structures, retaining paired concept/random results and all conformations within each structure. Random-encoder contrasts retain the same pairs across the five initializations. The reported percentile 95% intervals describe structure-sampling uncertainty conditional on those directions and initializations; they do not fully quantify uncertainty over alternative direction or pretraining draws. The original code also reports (1 + *k*)*/*(*B* + 1), where *k* counts bootstrap estimates at or below zero. We retain these as bootstrap tail fractions, not exact null-calibrated *p* values. Applying BH to them does not make them exact tests. A positive interval and a tail fraction are distinct summaries.

For MolCLR, aggregate accuracies alone do not determine the variance of a paired difference. The primary pair-level file was unavailable for this revision, so neither paired clustered intervals nor a primary hypothesis-test verdict are reported. A valid follow-up must retain concept/random outcomes together and account for repeated molecular structures across pairs. The two primary blocks should form a declared testing family. Approximate MDEs are planning quantities, not significance cutoffs or exclusion bounds (Appendix B).

### 3.4 Operational evidence ladder

We distinguish four levels: (1) recoverability not established against the specified decoding control; (2) linearly recoverable; (3) direction-specific intervention evidence in the fitted encoder–readout system; and (4) an additional pretraining-associated selectivity advantage against random initializations. The submitted paper called Levels 3 and 4 “causally accessible” and “pretraining-specific causal.” Here those names are explicitly restricted to this operational system, not to a native prediction head or a uniquely identified chemical mechanism.

For descriptive primary evidence, Level 3 requires a positive Δ_dir_ with its structure-clustered 95% interval above zero; Level 4 additionally requires the analogous result for Δ_enc_. Table 1 gives both. These are pointwise, conditional interval criteria, not a claim of multiplicity-controlled confirmation. The stricter empirical-direction and seed analyses are reported alongside them. MolCLR reaches Level 2, with higher-level evidence unresolved rather than refuted. Failure to establish one level is not proof that its underlying mechanism is absent.

**Table 1:** Primary Uni-Mol contrasts with structure-clustered, pointwise 95% percentile intervals. Each row satisfies the conditional descriptive Level 3 and Level 4 interval criteria. The 20-direction sensitivity family provides no corrected confirmation. Total/bond labels are joined by concept name to the saved analysis table.

| Block | Concept | $\Delta_{\text{dir}}$ [95% CI] | $\Delta_{\text{enc}}$ [95% CI] |
| --- | --- | --- | --- |
| 4 | total | 0.191 [0.177, 0.205] | 0.161 [0.149, 0.175] |
| 4 | bond | 0.189 [0.177, 0.202] | 0.166 [0.152, 0.178] |
| 8 | total | 0.430 [0.404, 0.455] | 0.269 [0.245, 0.294] |
| 8 | bond | 0.379 [0.357, 0.401] | 0.252 [0.230, 0.275] |
| 12 | total | 0.463 [0.437, 0.489] | 0.179 [0.152, 0.207] |
| 12 | bond | 0.419 [0.393, 0.444] | 0.172 [0.142, 0.201] |

## 4 Results

### 4.1 Uni-Mol: selectivity conditional on a fitted readout

At block 4, held-out *R*^2^ values are 0.783 for total distortion, 0.837 for bond distortion, and 0.859 for relative energy. At block 15 they are 0.703, 0.731, and 0.759, respectively. At block 12, total-distortion edits achieve 97.6% sign accuracy and Spearman *ρ* = 0.787; bond-distortion edits achieve 95.8% and *ρ* = 0.829. The six primary Δ_dir_ and Δ_enc_ intervals exclude zero (Table 1). All original one-sided bootstrap tail fractions equal the finite-resampling floor 1*/*2001 ≃ 0.0005; these are not presented as exact significance levels.

Total-distortion Δ_dir_ increases from 0.191 at block 4 to 0.430 at block 8 and 0.463 at block 12, while probe recovery declines. The corresponding Δ_enc_ values are 0.161, 0.269, and 0.179; bond-distortion values are 0.166, 0.252, and 0.172. The pretraining-associated component is therefore not monotonic.

These trends describe the tested edits and fitted readout. Without calibrated bypass or final-layer controls they cannot establish a greater contribution of subsequent encoder computation at later edit sites.

### 4.2 Concept–outcome alignment and an outcome-projection sensitivity analysis

The submitted manuscript identified alignment with the outcome readout as an unresolved explanation. Existing post-review artifacts now quantify the overlap (Table 2). Raw-coordinate concept directions have absolute cosine 0.472–0.939 with the same-block energy-probe direction. The leading correlation between orthonormal bases of the joint concept span and outcome span is 0.919–0.955. This is geometric subspace overlap, not an empirical correlation of predicted labels.

**Table 2:**
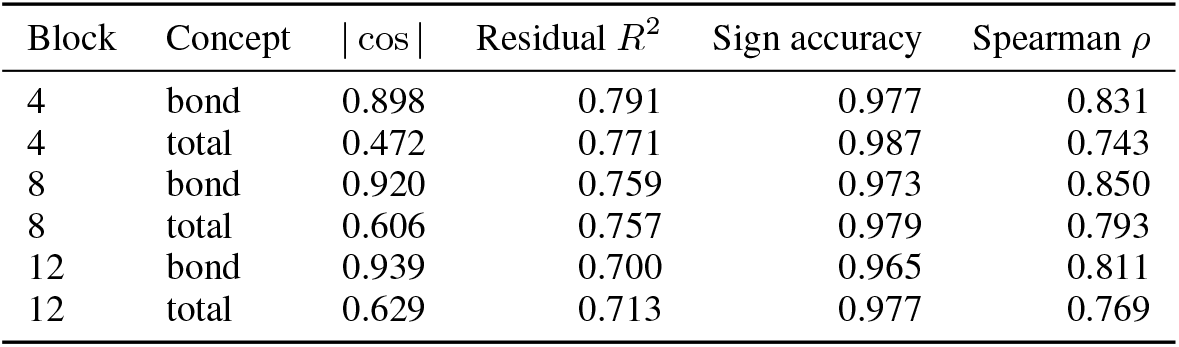
Post-review outcome-projection sensitivity analysis. The excluded direction is fitted at the edit block; the outcome readout is at block 15. These results do not constitute a matched bypass test.

The additional analysis projects out the same-block outcome span, refits the concept probe using the training/validation split, and reapplies the projector to the effective raw coefficient before normalization. Residual concept *R*^2^ is 0.700–0.791. Interventions along these residual directions retain sign accuracy of 0.965–0.987 and Spearman correlation of 0.743–0.850. We recomputed these metrics from all 186,600 saved pair-level rows (six cells of 31,100 pairs).

This analysis excludes alignment with the particular fitted same-block outcome direction by construction. It does not remove all information about energy, establish independence from correlated factors, or show that the remaining blocks are necessary. In particular, orthogonality to a same-block probe is not orthogonality to the effective sensitivity of the layer-15 readout. The saved projection run used CPU float32, whereas the primary run used GPU bfloat16, and it did not include a new matched random-direction family. We therefore treat persistence of the response as descriptive sensitivity evidence, not as a controlled improvement over the original edit or resolution of the downstream-computation alternative.

### 4.3 MolCLR: improved recovery, unresolved selective effect

At block 5, the pretrained concept probe has *R*^2^ = 0.385, compared with 0.299–0.313 for random encoders; outcome recovery is 0.439 versus 0.349–0.359. The mean concept-recovery advantage is 0.080 *R*^2^.

From the unrounded primary outputs, block-2 concept accuracy is 0.534515 and the random-direction mean is 0.505970, giving Δ_dir_ = −0.028545. At block 4 the values are 0.478545 and 0.493470, giving Δ_dir_ = −0.014925. Concept effect–outcome correlations are 0.055 and 0.018. These point estimates do not establish either a reliable selective effect or its absence. We withdraw the submitted exclusion claim based on the MDE; equivalence or exclusion requires an appropriate interval and a justified effect-size bound.

Local edit sensitivity analyses use uniform, top-node, and contribution-weighted selectors under pooled and Frobenius norm regimes. Their pretrained Δ_dir_ spans −0.018 to 0.021. Figure 4A instead plots Δ_enc_ at block 4, with uncertainty across random-encoder seeds; it is a different contrast and does not supply the missing primary paired interval. Three of its six pointwise seed intervals exclude zero, two in the negative direction and one in the positive direction. None of the twelve one-sided positive-advantage tests survives the saved BH correction across both blocks and all six selector/norm combinations. This does not imply equivalence.

**Figure 1:**
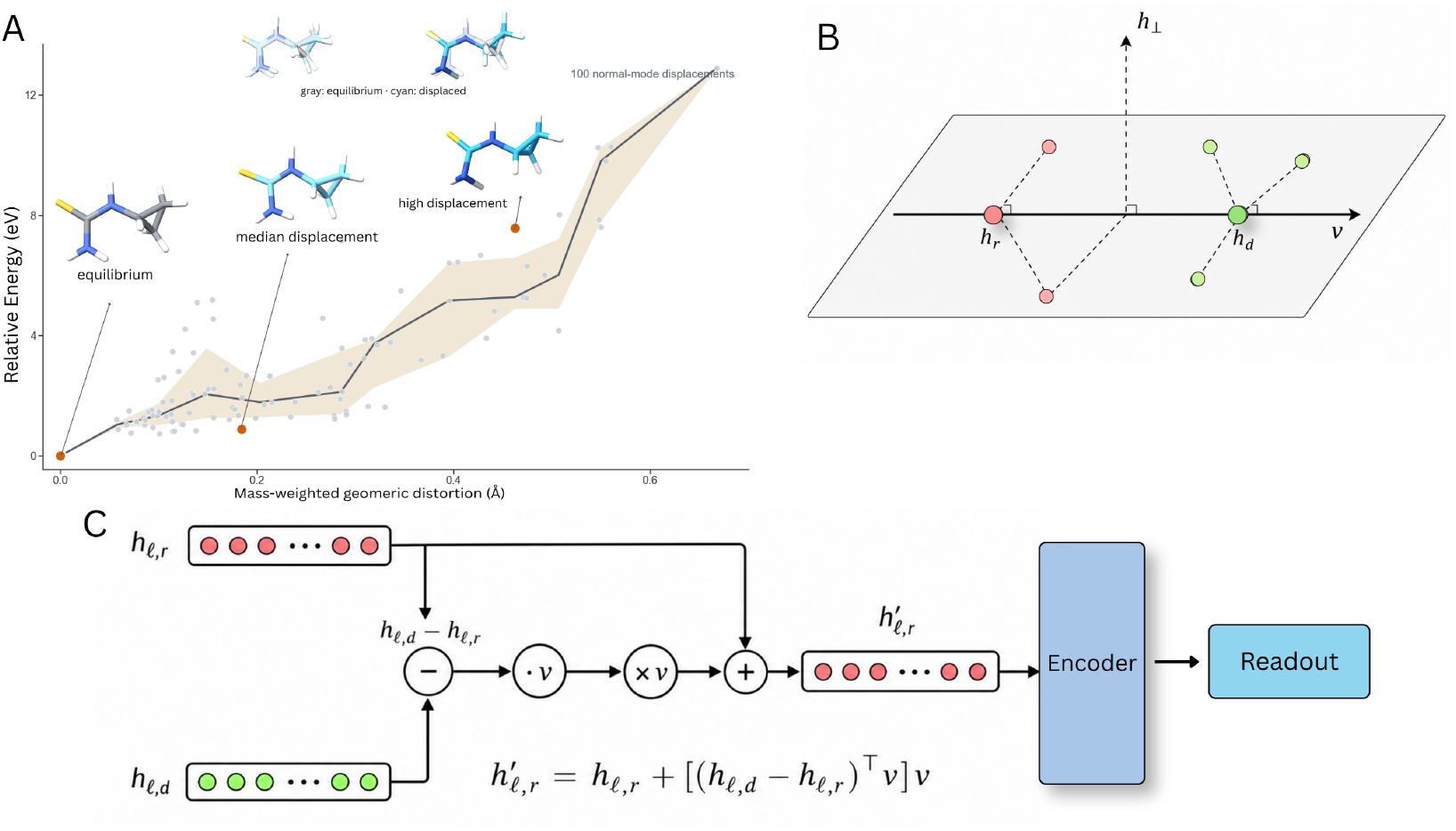
Intervening on a recovered direction in an encoder–readout system. A: one held-out QM7-X structure and 100 accepted normal-mode displacements; the line and band show median and 20–80% relative-energy values. B: projections onto a concept direction. C: donor-minus-recipient projection replacement, followed by remaining encoder computation and a fitted outcome readout. Hydrogens are omitted from the rendering only; all atoms enter the experiment.

**Figure 2:**
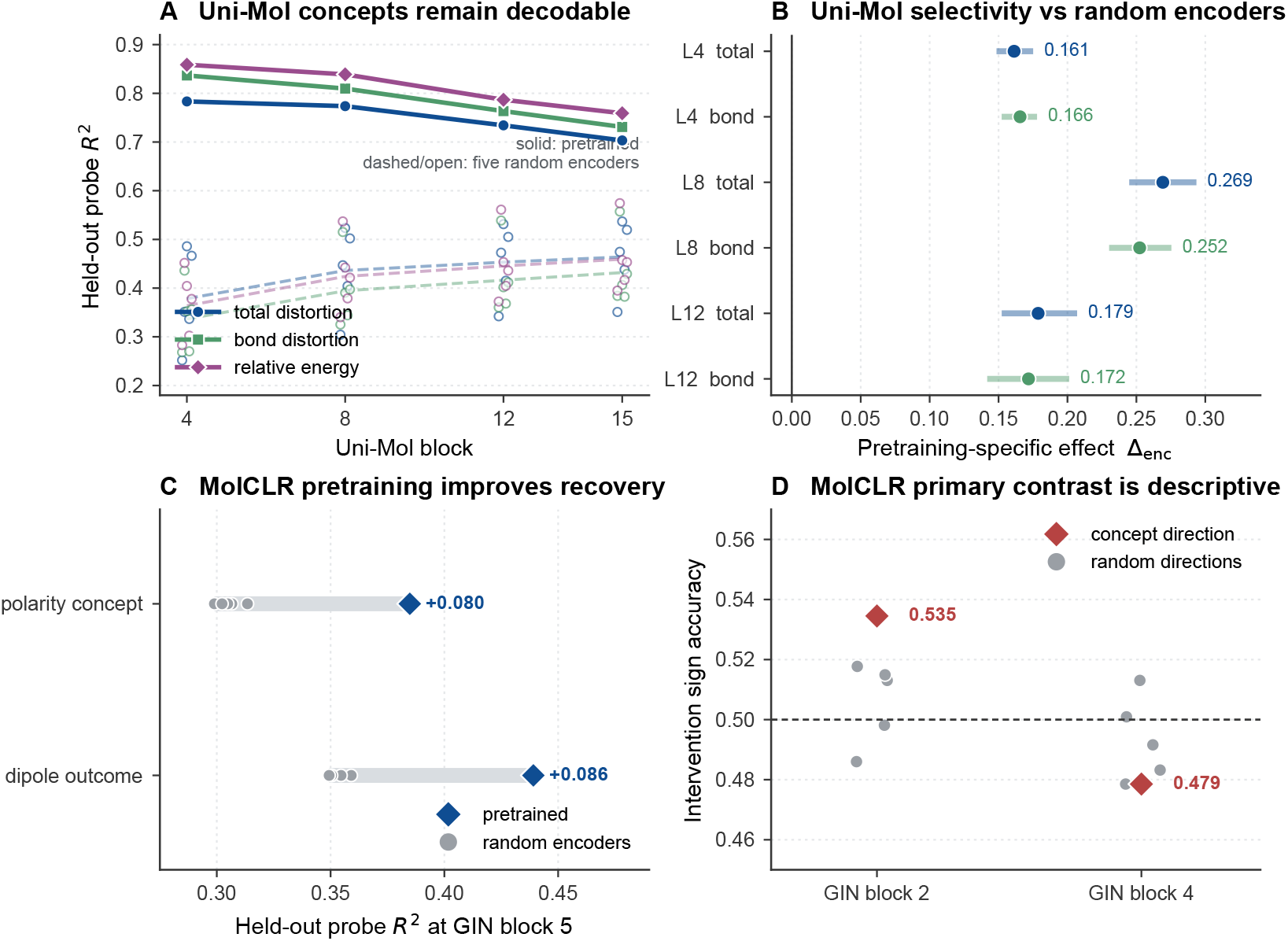
Recoverability and intervention selectivity are distinct measurements. A: Uni-Mol probe recovery; solid lines denote pretrained models, dashed lines and open markers summarize five random encoders. B: the six Δ_enc_ estimates with structure-clustered 95% intervals. C: MolCLR probe recovery at block 5. D: primary MolCLR concept accuracy and the five individual random-direction accuracies; their spread is not a confidence interval for Δ_dir_. The horizontal 0.5 line is a reference, not a required random-control outcome.

**Figure 3:**
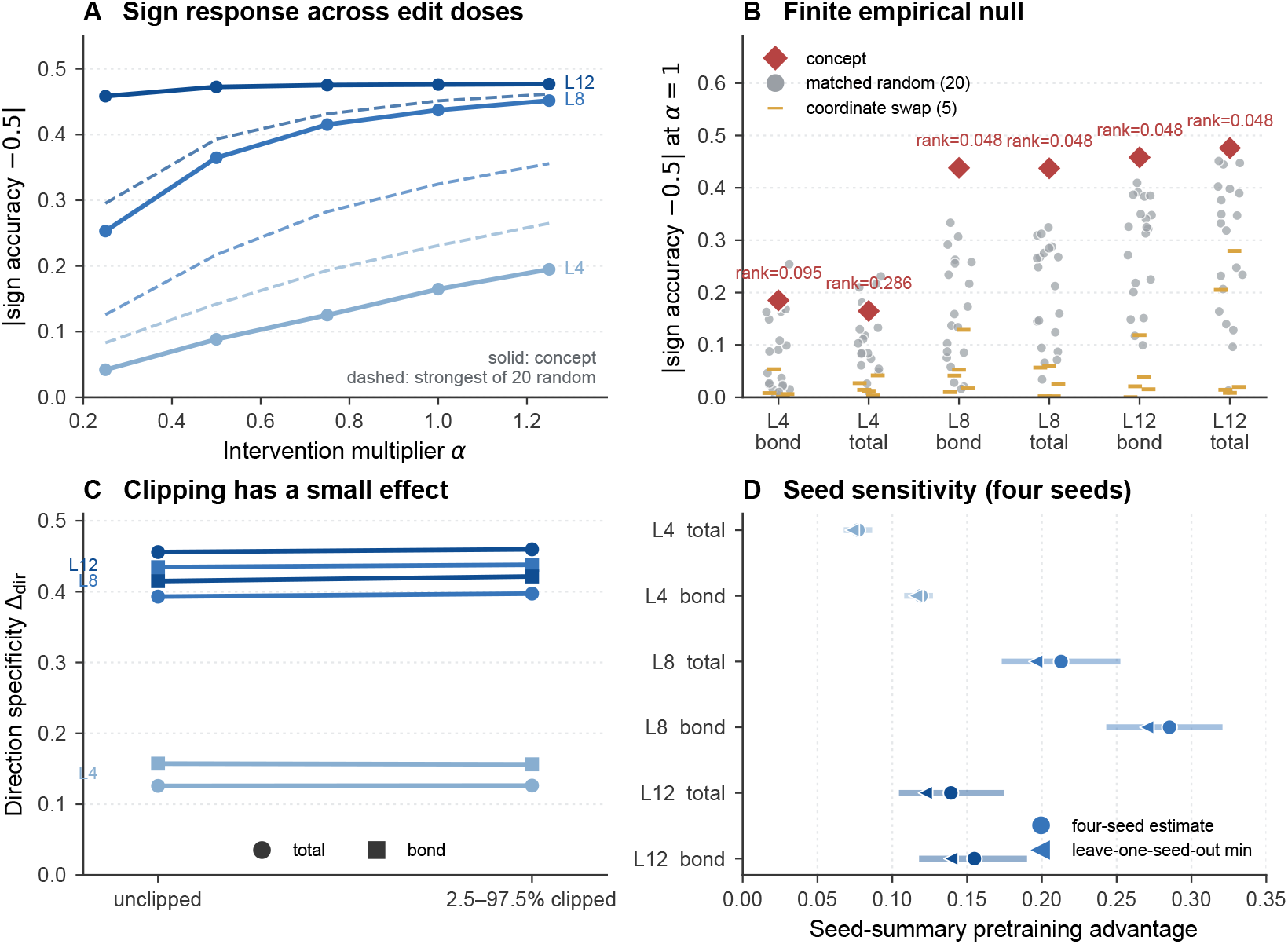
Uni-Mol sensitivity analyses use distinct estimands. A: absolute sign-accuracy deviation from 0.5 for total distortion across doses; dashed lines show the strongest of 20 random directions at each dose. B: full absolute-deviation distributions, with descriptive tail ranks; none reaches the corrected 0.05 threshold across six cells. C: raw-accuracy Δ_dir_ before and after clipping. D: Δ_enc_ sensitivity across four complete random-encoder seeds under 2.5–97.5% clipping; intervals resample seeds, not molecular structures.

**Figure 4:**
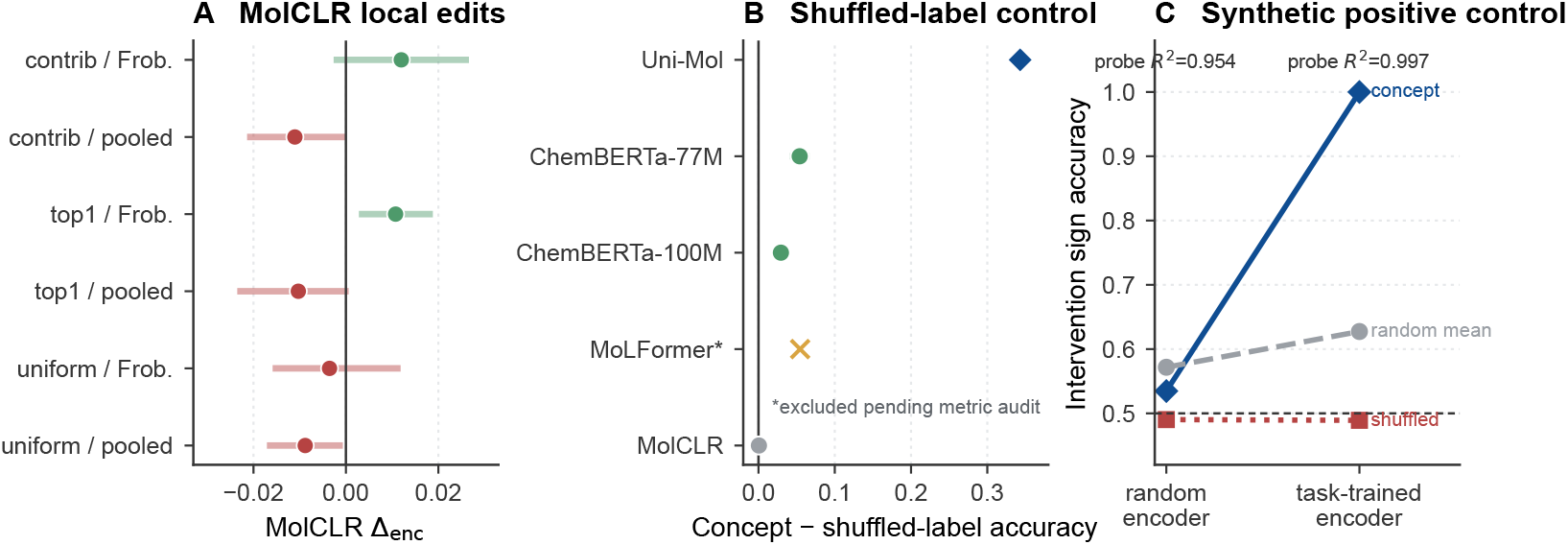
Additional controls bound interpretation. A: MolCLR block-4 Δ_enc_ for six selector/norm combinations, with seed-bootstrap intervals; none passes the one-sided FDR rule across the twelve comparisons at blocks 2 and 4. B: saved concept-minus-shuffled-label accuracy summaries, with MoLFormer marked as excluded from causal interpretation pending a metric audit. C: synthetic control averages over five seeds; random-encoder recoverability does not imply a privileged concept direction.

### 4.4 Robustness and negative controls

The 20-direction family matches each edit magnitude exactly. Median relative edit norms differ by at most 2.9 *×* 10^−6^ between concept and random edits across cells. This controls that aspect of edit size, but does not establish on-manifold states or rule out an aligned readout. Some random directions have substantial effect–outcome correlation (| *ρ*| up to 0.72 at block 12), motivating comparison with their full distribution.

The submitted empirical-null figure uses the two-sided descriptive statistic *T* (*v*) = |*A*(*v*) − 0.5 |, not the directional accuracy in Eq. 5. Its tail rank is

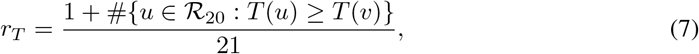

including ties. There are zero exceedances at blocks 8 and 12 for either concept; block-4 bond and total distortion have one and five, respectively. The ranks are thus 0.0476, 0.0952, and 0.2857. The minimum BH-adjusted rank across the six cells is 0.0714, so no cell reaches 0.05 under that sensitivity rule. These finite-direction ranks are not automatically exact permutation tests: the fitted concept direction and constrained random directions are not shown to be exchangeable under a null. The inference is descriptive even before multiplicity correction. A planned 200-direction extension is not included because completed outputs were unavailable.

Clipping changes the sensitivity-family Δ_dir_ by at most 0.004. Dose curves and their limitations are shown in Fig. 3. Coordinate swaps are not inert: for block-12 total distortion, mean sign accuracy is 0.594 and mean *ρ* is 0.101; the strongest swap achieves 0.779 and 0.324. Four-seed sign tests have a minimum one-sided value of 0.0625 and provide no 0.05-level confirmation. Positive seed-bootstrap intervals and leave-one-out estimates at blocks 8 and 12 are sensitivity evidence conditional on this small seed set.

## 5 Conclusion and limitations

Linear recovery and direction-specific intervention selectivity answer different questions. Uni-Mol supplies positive conditional contrasts within one distortion–energy family. The stricter direction and seed checks do not provide multiplicity-controlled confirmation, and the fitted-readout alignment alternative remains unresolved. MolCLR improves recoverability but its primary selective-effect uncertainty remains unquantified; it is not a precise negative result.

Neither case establishes that a native molecular prediction head uses the tested concept, that pretraining uniquely created the mechanism, or that an effect requires computation in the remaining blocks. Rank-one edits may miss nonlinear, distributed, or edge-state representations. Real donor projections can still create off-manifold combinations. The two distortion concepts share an outcome and are not independent replications. Uni-Mol and MolCLR differ in too many respects to attribute the contrast to modality or architecture.

A matched direct-readout/bypass or final-layer experiment, with comparable edit scales and readout calibration, is the clearest next test of downstream computation. Paired MolCLR intervals are needed before drawing an exclusion conclusion. Larger direction sets, additional concept–outcome families, and native-head evaluation would broaden the evidence. The reporting rule is to name the intervention, readout, control family, uncertainty, and unresolved alternatives alongside every mechanistic claim.

## A Construction and validation details

### QM7-X

For displaced conformations, the concepts are mass-weighted aligned RMS distortion and RMS bond-length distortion, computed without energy labels. Bonds are inferred from the optimized geometry. Total and bond distortion pass the fixed input-validity gate on 31,100 held-out interventions, with concept–outcome *ρ* = 0.808 and 0.956 and expected-sign rates of 1.0. Minimum nonbonded compression fails (*ρ* = 0.106, below the 0.20 gate). CLS states are extracted after blocks 4, 8, 12, and 15. Five reinitialized encoders use seeds 20260721–20260725.

### QM9

The Mulliken-derived dipole predicts DFT dipole change with *ρ* = 0.751 and expected-sign rate 0.940. Formula counts and 1,024-bit Morgan fingerprints (Rogers and Hahn, 2010) provide non-neural baselines. Uniform node edits use equal weights, top-node edits target the largest absolute concept contribution, and contribution-weighted edits use all contributions. Pooled and Frobenius regimes match aggregate or node-state norm, respectively.

### Metric checks

The revision’s runnable audit checks sign orientation on known positive and negative effects, exact-zero behavior, donor projection replacement, zero-dose identity, and invariance of the rank-one edit under *v* ↦ − *v*. It recomputes all six outcome-projection metrics from saved pair-level data and reproduces the empirical-direction ranks from saved metrics. These checks validate the stated conventions and available artifacts; they do not replace an end-to-end rerun of every encoder. In particular, no assumption of exactly 50% random-direction accuracy is used as an implementation test.

## B Power and uncertainty

The submitted MolCLR planning calculation uses *n* = 1072 and the binomial standard error at 0.5, 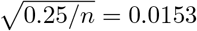. At two-sided *α* = 0.05 and 80% power, its approximate detectable shift from 0.5 is 0.0428, rounded to 0.043. An independence approximation for concept accuracy minus the mean of five random-direction accuracies gives 0.0469. Neither approximation accounts adequately for the paired indicators or possible reuse of structures, and neither is an observed-effect exclusion threshold. A future paired analysis should use the within-pair difference in correctness, retain all six indicators together during structure-level resampling, and account for structures shared across pairs. The two blocks should be analyzed under a declared multiplicity rule; seed-only uncertainty cannot substitute for pair-sampling uncertainty.

The primary Uni-Mol intervals resample structures, whereas the seed sensitivity intervals resample four complete random encoders using 20,000 draws. The original BH families for the primary bootstrap tail fractions contained six block–concept cells separately for Δ_dir_ and Δ_enc_. The empirical-direction family contains six cells; the MolCLR local-edit sign-test family contains twelve comparisons (two blocks, three selectors, two norm regimes). The Uni-Mol seed-sign family contains twelve block/concept/clipping rows and is descriptive because of incomplete cells and coarse seed resolution. Shared concepts and blocks create dependence. Positive association alone does not demonstrate the stronger dependence conditions required for a formal BH guarantee, another reason to keep descriptive summaries separate from confirmatory claims.

## C Synthetic validation and SMILES diagnostics

The synthetic control has 10,000 constructed pairs per seed and five seeds, reusing the same pairs across seeds. Its planning MDE from 0.5 is 0.014 per seed; 50,000 observations must not be treated as independent. A task-trained encoder reaches concept *R*^2^ = 0.997, outcome *R*^2^ = 0.9996, sign accuracy 1.0, and effect–outcome *ρ* = 0.854. Shuffled-label directions average 0.489. A random encoder reaches concept *R*^2^ = 0.954, but its concept accuracy of 0.535 is below its random-direction mean of 0.572. This controlled example demonstrates recoverability without a privileged fitted direction.

ChemBERTa-77M and ChemBERTa-100M (Chithrananda et al., 2020) recover polarity with *R*^2^ near 0.35, but their small intervention contrasts depend on serialization and do not establish a pretraining-associated advantage. We retain the description “fragile.” MoLFormer (Ross et al., 2022) has reported block-12 concept accuracy 0.169 versus a random-direction mean of 0.060; at block 6 both are about 0.060. A positive contrast between these values does not establish correct directional prediction. Below-0.5 values alone do not prove a metric error. The sign-convention/labeling concern remains unresolved, and the earlier causal-accessibility verdict remains withdrawn pending an end-to-end audit.

## D Frozen analyses, deviations, and missing cells

Blocks, concepts, splits, seeds, validity gates, and primary hypotheses were frozen before outcome inspection according to the study protocol. Four preregistered QM9 systems failed input-validity gates: small-ring count versus electronic energy (166 held-out pairs); graph diameter versus polarizability (*ρ* = 0.030); radius of gyration versus polarizability (*ρ* = 0.082); and conjugation versus gap (428 pairs, *ρ* = −0.167, expected-sign rate 0.341). Thresholds were not retuned. QM7-X was added by amendment with frozen hypotheses before its outcome inspection; the initial registration did not select QM7-X.

One Uni-Mol random seed completed the original 1–99% clipping grid, and the other four completed the reduced no-clipping and 2.5–97.5% grid. All five contribute to the primary analysis. Condition-matched sensitivity summaries use four complete seeds. The block-4 total-distortion/no-clipping row for seed 20260721 lacks the concept observation, so its derived random-encoder mean is undefined and the corresponding incomplete summary is excluded.

The primary block-4 total-distortion Δ_dir_ = 0.191 uses five original random directions. The post-primary value 0.126 uses 20 magnitude-matched directions; these are different contrasts. Outcome-projection results are post-review sensitivity analyses. A 200-direction analysis and matched bypass experiment are not reported as completed. This revision preserves the frozen results while correcting their labels and inferential interpretation.

## E Declarations and release status

### Data

QM9 and QM7-X are public datasets. The local revision package contains an audit of the available derived metrics and source hashes. A public code/results archive and its access conditions remain to be confirmed before release.

### Ethics

This study uses public molecular data and involves no human participants.

### Review status

Supervisor approval and confirmed authorship, affiliations, funding, acknowledgments, competing interests, and applicable AI-assistance disclosure are pending. This internal draft is not approved for submission.

## Notes

### Competing Interest Statement

The authors have declared no competing interest.

